# Investigating a possible role of *R3HCC1L* in embryonic development and ocular disease

**DOI:** 10.1101/2024.10.29.620958

**Authors:** Megan C. Fischer, Linda M. Reis, Sanaa Muheisen, Sarah E. Seese, Elena V. Semina

**Affiliations:** Department of Cell Biology, Neurobiology and Anatomy, Medical College of Wisconsin, 8701 Watertown Plank Road, Milwaukee, WI, 53226, USA; Department of Ophthalmology and Visual Sciences, Medical College of Wisconsin, 8701 Watertown Plank Road, Milwaukee, WI, 53226, USA; Department of Pediatrics and Children’s Research Institute, Medical College of Wisconsin and Children’s Wisconsin, 8701 Watertown Plank Road, Milwaukee, WI, 53226, USA

## Abstract

Peters anomaly (PA) is an anterior segment ocular disorder with wide phenotypic variability and genetic heterogeneity. Here we report a family consisting of a male with a diagnosis of syndromic PA and his unaffected parents, with no causative variants identified in known developmental ocular genes. Exome sequencing analysis identified compound heterozygous missense variants, c. 1022A>T p.(Asp341Val) and c.1457T>A p.(Phe486Tyr), in *R3HCC1L*. Both variants are ultra-rare in control populations and have CADD scores of 21.9 and 23.3, respectively, suggesting possible deleterious effects. The *R3HCC1L* transcript variants encode three different protein isoforms all sharing two conserved C-terminal domains, an RNA recognition motif (RRM) and a coiled-coil domain (CCD), and likely represent an RNA-binding protein involved in post-transcriptional gene regulation; the identified patient variants are located within the N-terminal part of the protein shared by two of the three protein isoforms, upstream of the RRM and CCD domains. To investigate the possible role of *R3HCC1L* in embryonic development, the single zebrafish ortholog of *R3HCC1L, r3hcc1l*, was examined for its expression and function. *In situ* expression studies showed that zebrafish *r3hcc1l* is expressed in the developing lens, cornea, retina and hyaloid vasculature, supporting its possible role in ocular development in vertebrates. CRISPR-Cas9 gene editing was used to generate a zebrafish line with a 4-bp deletion in *r3hcc1l*, c.623-626del, that is predicted to result in a nonsense-mediated decay and, if expressed, a nonfunctional truncated protein (p.Thr208fs*39) lacking 80% of the RRM and the entire CCD. The resultant *r3hcc1l*^*c*.*623-626del*^ heterozygous and homozygous animals did not show any visible structural abnormalities in the eye or any other systems, with normal survival of all genotypes to adulthood, providing no support for its possible role in the congenital phenotype of interest. However, the function of *r3hcc1l* may not be completely conserved with human *R3HCC1L*, and/or zebrafish may have compensatory mechanisms that are not present in humans. In addition, the engineered zebrafish variant disrupts the most conserved C-terminal region of R3HCC1L/r3hcc1l shared by all protein isoforms and likely leads to a complete loss-of-function of this gene, which may be different from the disease mechanism associated with the specific missense alleles identified in the patient. Finally, while it is important to consider the possible limitations of animal models, it is also necessary to highlight that the identified *R3HCC1L* variants may not have any role in the phenotype observed in this single patient. Identification of new *R3HCC1L* variants of interest in families affected with PA or other congenital phenotypes, if successful, will provide further support for the possible developmental function of this gene.

## Introduction

Peters anomaly (PA) is a rare ocular phenotype that falls under the spectrum of anterior segment disorders (ASD). It is characterized by central corneal opacity, corneolenticular and/or iridolenticular adhesions, and defects in the posterior corneal layers. PA can be isolated or can be associated with other ocular and systemic features. A wide array of systemic features has been reported including central nervous system abnormalities, congenital heart defects, dysmorphic facial features, growth retardation, syndactyly, developmental delay, hearing loss, cleft lip/palate, and skeletal abnormalities (Elbaz et al., 2022; Weh, Reis, Tyler, et al., 2014). Aligning with this phenotypic variability, PA is genetically heterogeneous. Most cases are sporadic and without an identified genetic cause although inherited forms have been documented with both autosomal dominant (*FOXC1, PAX6, PITX2*) and recessive (*B3GLCT, CYP1B1*) inheritance patterns (Chesneau et al., 2022; Reis et al., 2024; Weh, Reis, Happ, et al., 2014). Peters plus syndrome (PTRPLS) [MIM: 261540] which presents specifically with PA/ASD, short stature, dysmorphic facial features, and brachydactyly with variable other features is caused by autosomal recessive variants in *B3GLCT* (Weh, Reis, Tyler, et al., 2014). While progress has been made in finding genetic causes, many cases of PA remain undiagnosed. Additionally, many of the genes that have been associated with PA are more commonly associated with other ocular phenotypes, such as *FOXC1* and *PITX2* which typically cause Axenfeld-Rieger Syndrome (Reis et al., 2023). Continued investigation of underlying genetic causes is necessary to better understand the spectrum of phenotypes in ASD and PA and to create better classification systems. It has also been shown that the severity of ocular phenotypes in PA is not predictive of systemic involvement (Elbaz et al., 2022) and thus, classification by genetic cause may allow for better risk assessment for potentially life-threatening systemic abnormalities and help direct clinical management.

The incorporation of next generation sequencing (NGS) techniques into the clinic has been essential to the pursuit of a precise molecular diagnosis across ocular disorders. However, NGS still has many limitations and in PA many cases remain unexplained. Careful re-analysis of these unexplained cases is necessary to further our understanding of the underlying genetic causes. While this reanalysis must consider many explanations for a lack of diagnosis, including poor coverage or variants in unidentified regulatory regions of known disease genes, novel gene discovery is of particular interest for PA. Evaluating variants in novel genes requires moving beyond the predictive bioinformatics tools and into model systems to provide further evidence to support a role in development and disease.

*Danio rerio*, commonly known as zebrafish, are a popular model for the study of ocular development and disease. Ocular development in zebrafish closely resembles that of humans, with contributions from the same three embryological tissues: neuroectoderm, surface ectoderm, and mesenchyme (Richardson et al., 2017). Specifically, the development of the anterior segment is largely conserved with the lens placode budding off from the surface ectoderm, which develops into the cornea after complete lens detachment. This is important because PA arises from failure of the lens to separate completely from the overlying surface ectoderm in early development (Matsubara et al., 2001). It is worth noting that there is a key difference between humans and zebrafish in these developmental steps. In humans, the lens placode invaginates forming a vesicle at its center known as the lens pit, whereas in zebrafish the lens placode forms a solid mass of cells which detach by apoptosis (Greiling et al., 2010). Regardless, there are established genetic models in zebrafish that demonstrate disruption in the development of cornea and/or lens following mutation of orthologs of known human PA genes including *pitx2* (*PITX2), foxc1a/b* (*FOXC1)*, and *pax6a/b* (*PAX6)* (Ferre-Fernandez et al., 2020; Hendee et al., 2018; Takamiya et al., 2020). In addition to having similar developmental processes to interrogate, zebrafish also have a great amount of genetic overlap, with about 70% of human genes having at least one zebrafish ortholog (Howe et al., 2013). This makes them a useful tool for genetic studies, which is further complemented by their ability to be genetically modified by a variety of tools and their production of large clutch sizes with short generation times.

Here we present an unexplained case of syndromic PA, in which we identified compound heterozygous missense variants in a gene of interest, *R3HCC1L*, as well as functional studies of its zebrafish ortholog, *r3hcc1l*, using CRISPR-Cas9-mediated genome editing.

## Methods

### Human Subjects and Exome Sequencing

This human study was approved by the Institutional Review Boards of the Medical College of Wisconsin and University of Iowa, with written informed consent obtained for every participant. Genomic DNA for the proband and both parents was submitted for exome sequencing to Psomagen (Rockville, MD). Variant locations were annotated to human Genome Build hg19. Trio analysis was performed using VarSeq (Golden Helix, Bozeman, MT) and variants of interest were annotated using the following tools: gnomAD v3.1.2 and gnomAD v4.1.0 (Chen et al., 2024), CADD GRCh37/hg19 (Kircher et al., 2014), and REVEL (Ioannidis et al., 2016). Samples were first analyzed for variants in genes known to be involved in PA or other ASD (see Reis et al., 2024 for the gene list), then variants in novel genes were evaluated according to inheritance pattern (homozygous, compound heterozygous, de novo, and X-linked). Samples were also evaluated for copy number variants using the VarSeq CNV Caller (VS-CNV) (GoldenHelix). Variants of interest were evaluated for splicing effects using the SpliceAI Lookup tool from the BROAD Institute (https://spliceailookup.broadinstitute.org/) giving scores for SpliceAI (Jaganathan et al., 2019) and Pangolin (Zeng & Li, 2022). Evaluation of variants was completed in accordance with the American College of Medical Genetics/Association for Molecular Pathology criteria (Richards et al., 2015).

### Expression Studies

*In Situ* hybridization was performed for *r3hcc1l* as previously described (Seese et al., 2021) on zebrafish whole embryos and head sections at 24 and 48 hours post-fertilization (hpf) using an RNAscope Probe (Cat No. 1209671-C3; Advanced Cell Diagnostics, Inc.).

### Multiple Sequence Alignment

Multiple sequence alignment between human *R3HCC1L* (NP_001337944), mouse *R3hcc1l* (NP_803415), and zebrafish *r3hcc1l* (XP_690294.1) was performed using EMBL-EBI Clustal Omega MSA tool (CLUSTAL 0(1.2.4)) (Madeira et al., 2024).

### Generation of r3hcc1l mutant line

Zebrafish were maintained as previously described (Seese et al., 2021). All zebrafish work was approved by the Institutional Animal Care and Use Committee at the Medical College of Wisconsin. CRISPR/Cas9 genome editing was used to generate a zebrafish line carrying a frameshift mutation in *r3hcc1l*. A crRNA (targeted to exon 3 of *r3hcc1l*) [5’-CCCAACCGAATTCAAGACGG] was annealed with tracrRNA in Nuclease-Free Duplex Buffer (Integrated DNA Technologies (IDT), Coralville, IA). The resultant gRNA was added to an injection mixture containing 2.5 μg Cas9 endonuclease in Cas9 working buffer (20 mM HEPES; 150mK KCl, pH7.5) and 0.1% phenol red. Approximately 4 nL of the mixture was injected into 1-4 cell stage embryos using the Eppendorf FemtoJet 4 Microinjector (Thermo Fisher Scientific, Waltham, MA). Injected embryos were raised to maturity (3 months) and out-crossed to the ZIRC WT-AB background to generate F1 embryos with a pool of mutations. Two founder pairs (F0) with a 4bp deletion (c.623-626del) were selected. For genotyping, DNA was extracted from whole embryos or adult fin clippings digested in NaOH and neutralized with Tris-HCl.

DNA was then amplified using *r3hcc1l*-specific primers flanking the target site (5’-GGAAGGACGCAAGAAGCAGT (ex3), 5’-GCACCGACACATTAATCACCC (int3-4), 531 nt) and sequenced (Functional Biosciences, Madison, WI).

Founder pairs (+/-) were bred and resulting F1 embryos were maintained in plates in E2 media in an incubator at 28.5C up to 5 dpf. Embryos were observed using a Zeiss Stemi 2000 Stereo Microscope (Carl Zeiss Inc., Thornwood, NY) from 0-5 dpf. Fish were then grown to adulthood (2-3 months) and assessed for gross morphology as well as genotype-specific survival effects using the Chi-squared statistical test in Microsoft Excel (Microsoft Corporation, Redmond, WA). Adult fish were imaged using a Zeiss Discovery V12 Microscope (Carl Zeiss Inc.). Additional breedings were then completed between F1 adult fish including homozygous (-/-) female with heterozygous (+/-) male and homozygous (-/-) with homozygous (-/-). These F2 embryos were maintained and observed as described above.

### Evaluation of zebrafish transcripts

Zebrafish F2 embryos (homozygous for *r3hcc1l*^*c*.*623-626del*^ and wild type) were collected in TRIzol (Thermo Fisher Scientific) at 4 dpf. RNA was extracted and purified using the RNA Clean & Concentrator-5 Kit (Zymo Research, Irvine, CA). 1.5 ug RNA was converted to cDNA using the SuperScript III RT-PCR System (Thermo Fisher Scientific) using random hexamer primers. DNA was then amplified with a first step of 3 mins at 94°C followed by 20 seconds at 94°C, 30 seconds at 60°C and 30 seconds at 72°C for 40 cycles, and then a final step of 7 minutes at 72°C using *r3hcc1l* primers flanking the target site (5’-GGAAGGACGCAAGAAGCAGT (ex3), 5’-GGAGAACAGACCAAGAGCGT (ex4), 235 nt) and sequenced (Functional Biosciences). DNA amplification was repeated with a shorter program, using 25 cycles, with the same primers for *r3hcc1l* and additional primers for *eef1a1l1* (5’-CTGGAGGCCAGCTCAAACAT (ex3-4), 5’-ATCAAGAAGAGTAGTACCGCTAGCATTAC (ex4), 87 nt). PCR products were resolved using agarose gel electrophoresis.

## Results

### Identification of R3HCC1L variants

A male with a diagnosis of syndromic PA with no further clinical details available and his unaffected parents were enrolled into the genetic study (Figure 1A). Trio WES analysis was negative for any pathogenic or likely pathogenic variants in known genes associated with PA or ASD. Initial filtering by allele frequency in the general population (AF<0.01; gnomAD v3.1.2), inheritance pattern, and read depth (RD>5) identified 13 homozygous, 21 de novo, 12 X-linked and 141 compound heterozygous variants for more careful analysis. Variants were further filtered by CADD>15, and compound heterozygous variants were restricted to variants in genes where at least one of the two was a coding variant. Only four compound heterozygous variants in two genes (*TAS1R3, R3HCC1L*) met these criteria and were further evaluated for allele frequency and gene function. The *TAS1R3* variants (NM_152228.3:c.465G>C, NM_152228.3:c.530G>A) were determined to be of lower interest due to their high allele counts in gnomAD v4.1.0 (914/1612216 with 10 homozygotes; 163/1559492 with 1 homozygotes) and the known gene function of *TAS1R3* which encodes a taste receptor (Sanematsu et al., 2014). The variants in *R3HCC1L* (NM_001351015.2= MANE transcript) were prioritized for further consideration. These compound heterozygous missense variants, c.1022A>T p.(Asp341Val) (paternal) and c.1457T>A p.(Phe486Tyr) (maternal) were present in the proband (54/107 and 16/41 reads respectively), each inherited from one unaffected parent (father (47/82 reads) and mother (25/41 reads), correspondingly) (Figure 1B). The paternal variant is ultra-rare (3/1613962 in gnomAD v4.1.0 with 0 homozygotes) and has some predictive evidence of being deleterious with a CADD score of 21.9 (but a REVEL score of 0.101). It is also reported in ClinVar (VCV003150477.1) as a variant of uncertain significance, although no other details are available. The maternal variant is also ultra-rare (4/161400 in gnomAD v4.1.0 with 0 homozygotes) and has some predictive evidence of being deleterious with a CADD score of 23.3 (but a REVEL of 0.037). Both CADD and REVEL are computational tools for scoring the deleteriousness of single nucleotide variants with a CADD score >20 indicating a variant is in the top 1% of deleterious variants in the human genome (Kircher et al., 2014) and REVEL scores ranging from 0-1 with a higher score indicating a higher probability of pathogenicity (Ioannidis et al., 2016). Neither variant had a predicted effect on splicing by SpliceAI or Pangolin.

**Figure 1:**
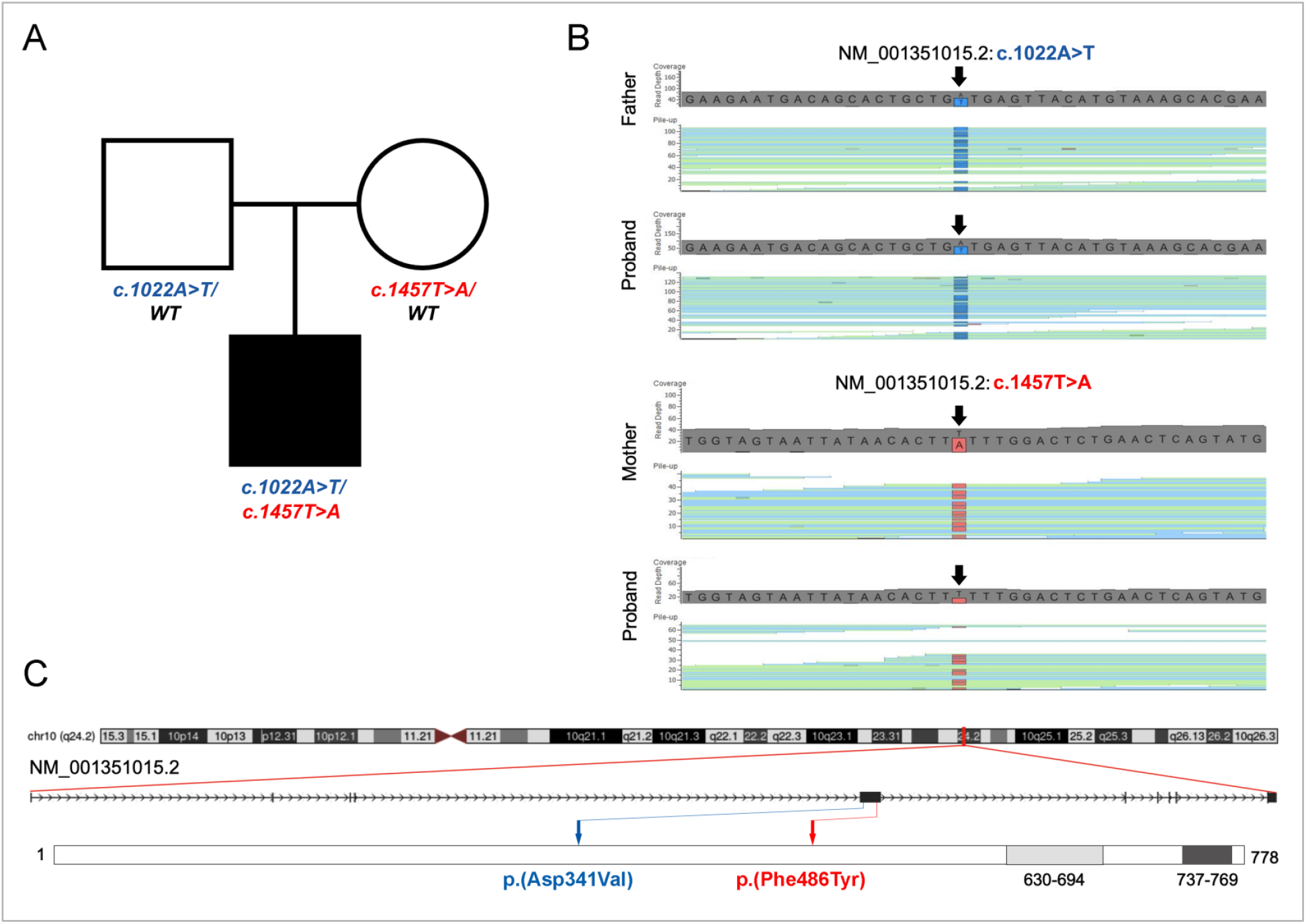
(A) Pedigree of the family; solid symbol indicates affected; WT indicates wild type sequence at the location of the variant. (B) BAM alignment showing presence of both variants in the proband and the c.1022A>T and c.1457T>A variants in the father and mother respectively. (C) *R3HCC1L* genomic location (top), MANE transcript (NM_001351015.2) (middle), and protein domain (bottom) structures with the identified variants indicated with blue (c.1022A>T) and red (c.1457T>A) arrows. RNA recognition motif (RRM) and coiled-coil domain (CCD) are shown in as light and dark grey boxes, correspondingly.

Both variants are located in exon 5 of *R3HCC1L* (out of 10 in the MANE transcript) and are upstream of the two annotated domains of this protein, an RNA recognition motif (RRM) (aa 630-694) and a coiled-coil domain (CCD) (aa 737-769) (Figure 1C). A number of different transcripts have been reported for this gene with 13 unique transcripts and more than 70 predicted transcripts included in NCBI and Ensembl; the main differences in these transcripts are in the inclusion of various 5’ non-coding exons; transcript variant 11 was designated as MANE transcript. In terms of protein variants, there are 3 different proteins encoded by the 13 unique *R3HCC1L* transcripts: transcript variant 1 encodes isoform 1 of 792 amino acids (aa), transcript variants 2-4 and 6-13 encode isoform 2 of 778 aa, and transcript variant 5 encodes the shortest isoform, isoform 3 of 184 aa; all proteins variants share the same C-terminal sequence containing RRM and CCD domains (Figure 2). GTEX human expression data shows that the most highly expressed transcripts are those that encode protein isoforms 2 and 3 with expression detected across all tissues (of note, eye tissues were not tested and the data is limited to only a subset of *R3HCC1L* transcript variants) (GTEx Analysis Release V8 (dbGaP Accession phs000424.v8.p2)).

**Figure 2:**
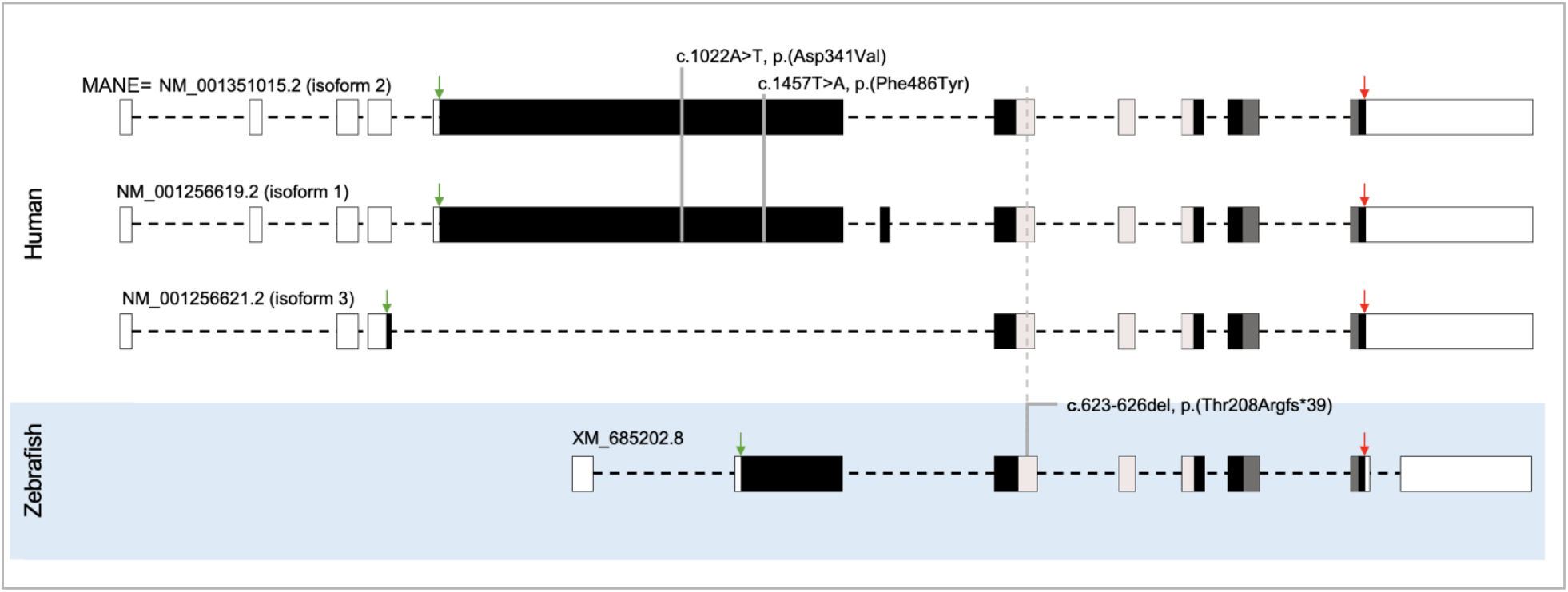
Partially scaled diagrams (exon sizes are in scale; intron sizes are not) denoting the human (top) and zebrafish (bottom) *R3HCC1L/r3hcc1l* transcript variants. The MANE transcript NM_001351015.2 encodes isoform 2 (top), NM_001256619.2-isoform 1 (the longest isoform; middle), and NM_001256621.2-isoform 3 (the shortest isoform; bottom). Patient variants are shown in the first coding exon of isoforms 1 and 2 (not present in isoform 3). Location of the zebrafish variant is indicated in the zebrafish transcript and corresponding locations in the human transcripts are shown with a dashed line. Translation start (green arrow) and termination (red arrow) sites, as well as regions encoding RRM (light grey) and CCD (dark grey) domains are indicated.

To further assess a possible role for *R3HCC1L* in human disease, various publicly available sources were evaluated for information about this gene. *R3HCC1L* does not show any involvement in developmental human disease; one study identified a possible association between *R3HCC1L* with greater pneumonia susceptibility and severity in a genome-wide association study, replicated in two independent datasets (Chen et al., 2021). It has a predicted role in nucleotide and/or protein binding based on the presence of an RRM and a CCD (Figure 1C) and thus R3HCC1L most likely functions as an RNA-binding protein (RBP) involved in post-transcriptional gene regulation (UniProt, 2023; Wang et al., 2023). *R3HCC1L* is also predicted to co-localize with the exon-exon junction complex (EJC) (Alliance of Genome Resources, 2019). Previous studies identified RBP-encoding genes as well as factors associated with the EJC as causes for various developmental phenotypes including corneal/lens anomalies (Aryal et al., 2020; Dash et al., 2015; Lachke et al., 2011; Richieri-Costa & Pereira, 1992), providing further support for this gene of interest. In terms of expression, *R3HCC1L* is known to have broad distribution across adult tissues, similar to other RBP genes, while its expression in early development is not well-characterized ((Thul et al., 2017); Human Protein Atlas proteinatlas.org). Similarly, mouse *R3hcc1l* is expressed broadly across all tested tissue types ((Lattin et al., 2008; Wu et al., 2009); biogps.org), including cornea, lens and other ocular structures, with the lens having the highest expression (greater than the median) over all other ocular tissues based on one probe set. The International Mouse Phenotyping Consortium ((Groza et al., 2023); www.mousephenotype.org) provides data for an unpublished *R3hcc1l* gene deletion mouse model (*R3hcc1l*^*tm1b*^*(KOMP)*^*Wtsi*^). Both male and female homozygous mice had significant phenotypes of decreased mean corpuscular hemoglobin and mean corpuscular volume. In the female mice only, a significant phenotype of the persistence of the hyaloid vascular system in the eye was observed (female mice: both eyes 4/8, one eye 3/8, and not present in 1/8; male mice: not present in 8/8). Additionally, 3/8 homozygous female mice had corneal opacity in one eye, although this was not observed in male mice and did not reach the threshold for significance. Both the persistence of the hyaloid vasculature and corneal opacity are developmental eye phenotypes, with the latter showing a particular overlap with the corneal opacity observed in our patient. Additionally, the broader phenotype of persistent fetal vasculature (PFV) is an important differential diagnosis in PA, as some cases of congenital corneal opacity can develop secondary to PFV, especially when corneal opacity develops shortly after birth (Franco et al., 2024; Huang et al., 2023). In summary, the available functional, expression, and animal model information for *R3HCC1L* supported its possible role in syndromic corneal opacity, though the data is very limited due to paucity of previous studies of this gene.

### Investigating zebrafish ortholog r3hcc1l

To investigate the possible role of *R3HCC1L* in embryonic development, we performed studies of its zebrafish ortholog (Figure 3). Currently, a single zebrafish *r3hcc1l* gene with a distinct transcript (XM_685202.9) has been identified; the zebrafish gene demonstrates conserved genomic structure and nucleotide sequence with the human gene in its 3’ region (including the five final coding exons) and a more divergent 5’ sequence, encoding a protein of 338 amino acids. Similarly, comparison with the human protein demonstrates diversity in the N-terminal region while the C-terminal region shows moderate conservation (45.27% identity overall) and a higher identity within the two annotated domains: RRM (aa 195-259) and CCD (aa 302-334) (72.31% and 68.75% level of conservation, respectively). The human variants identified in the affected individual, described above, are located in the region of no/poor conservation between the two orthologs. Because of this, it was not possible to interrogate the exact human variants in zebrafish, however, considering that recessive missense variants in humans are typically associated with a loss-of-function mechanism (Turner et al., 2015), investigation of a loss-of-function model in zebrafish was pursued.

**Figure 3:**
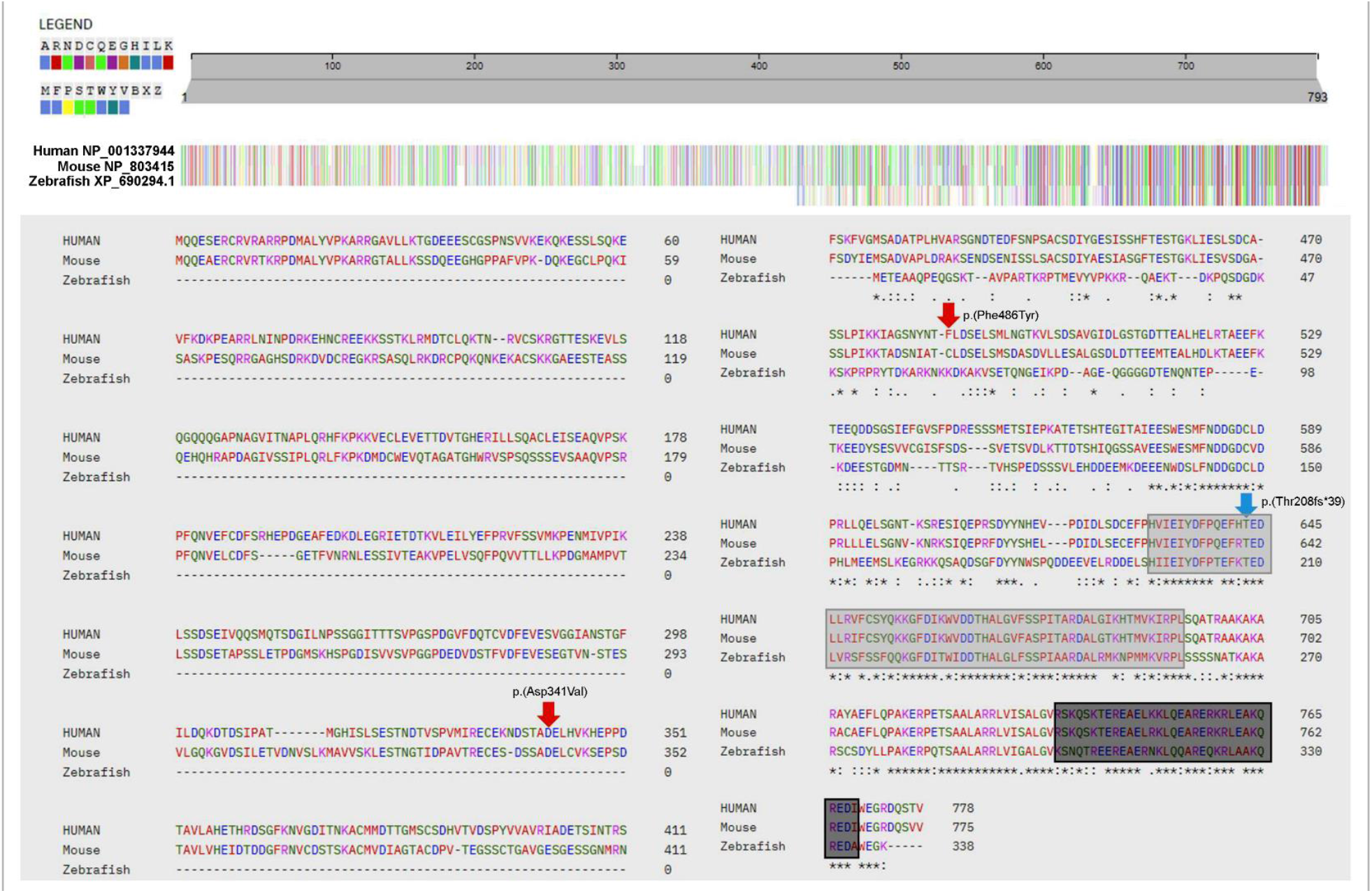
A multiple sequence alignment (MSA) of human, mouse, and zebrafish R3HCC1L proteins. MSA of the full protein (top) and at the amino acid level (bottom) are shown. Amino acids are colored following the Clustal2 scheme (legend in upper left). Red arrows indicate the locations of the two patient variants, blue arrow indicates the location of the zebrafish variant. Light grey box indicates the RRM, dark grey box denotes the CCD. Human R3HCC1L isoform 2 (NP_001337944), mouse R3hcc1l (NP_803415), and zebrafish r3hcc1l (XP_690294.1).

To further investigate the zebrafish *r3hcc1l* and whether it may contribute to ocular development, expression was investigated in early developmental stages. *In situ* expression studies showed that *r3hcc1l* is expressed in the zebrafish embryo (24-48 hpf (hours post-fertilization); Figure 4). More specifically, expression was seen in the developing brain and throughout the developing lens, cornea, and retina at 24 hpf (Figure 4A-D) with the expression continuing at all sites as well as hyaloid vasculature at 48 hpf (Figure 4E-H). This expression data supports a potential role for *r3hcc1l* in ocular development.

**Figure 4:**
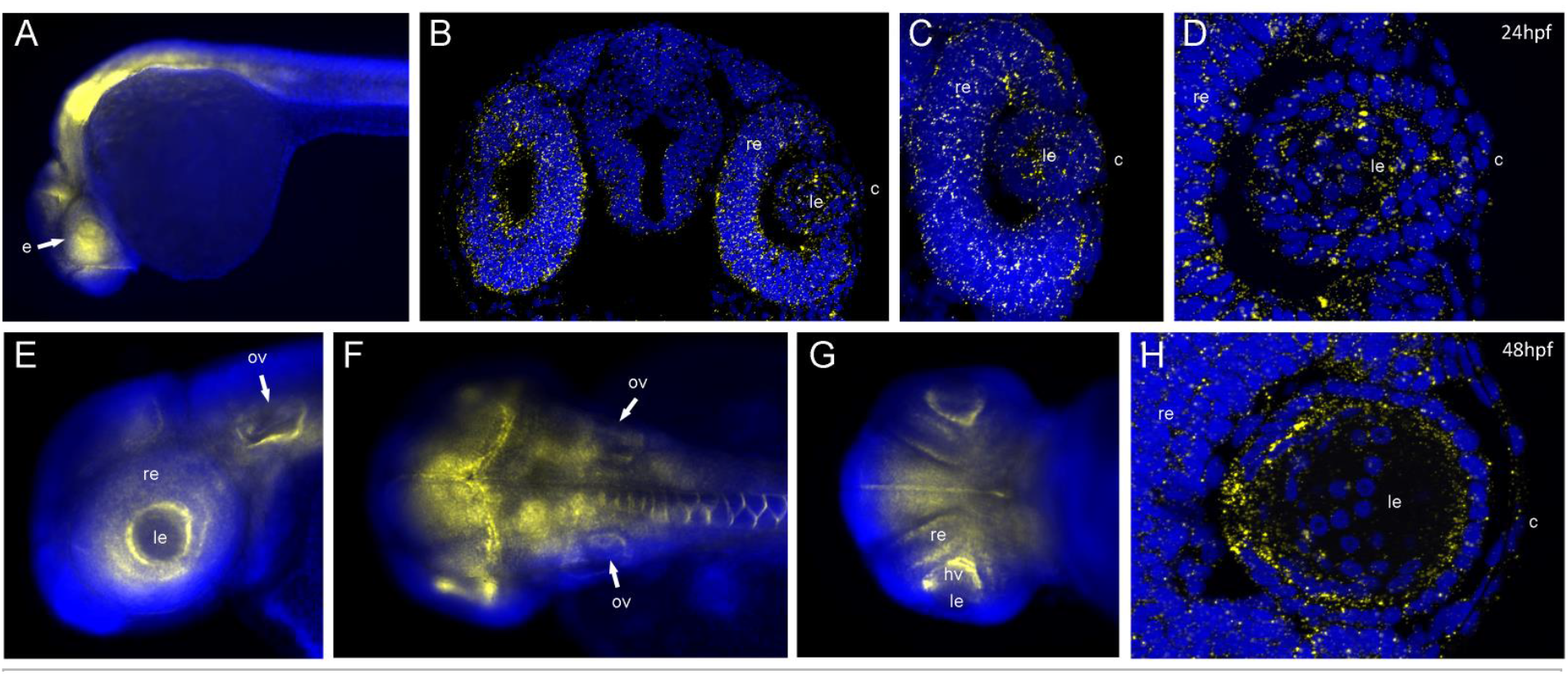
Expression pattern of *r3hcc1l* in zebrafish embryos. 24-(A-D) and 48-hpf (E-H) whole mount embryos (A, E-G) and transverse sections (B-D and H) are shown. Please note broad expression of *r3hcc1l* (yellow) in the developing brain, otic vesicles and the developing lens, cornea, and retina within the eye. DAPI (blue) was used for a nuclear counterstain. e- eye; c- cornea; le- lens; ov- otic vesicle; re-retina, hv- hayloid vasculature.

### Zebrafish mutant r3hcc1l have normal development and viability

CRISPR-Cas9 gene editing was used to generate a zebrafish line with a 4 bp deletion in *r3hcc1l (*c.623-626del; Figure 5A) that is predicted to lead to a frameshift and premature stop codon (p.Thr208fs*39). This mutation is 338 nucleotides (nt) upstream of the last exon-exon boundary and is thus predicted to be subject to nonsense mediated decay and lead to loss-of-function. If a protein product is made it would be significantly truncated (246 aa) and lack 80% of its RRM and the entire downstream CCD, and thus be unlikely to retain its function. To investigate mutant transcript stability, total RNA was extracted from homozygous *r3hcc1l*^c.623-626del^ and wild-type embryos (n=20 embryos at 4 dpf (days post-fertilization)) and *r3hcc1l* transcript was analyzed by semiquantitative RT-PCR with exon 3 and exon 4-specific forward and reverse primers, respectively (Figure 5B). Amplification with primers for eukaryotic translation elongation factor 1 alpha 1 like 1 (*eef1a1l1*) was completed in parallel as a house-keeping gene expression control. A visible reduction in mutant transcript level was detected for *r3hcc1l* but not for *eef1a1l1*, supporting partial NMD.

**Figure 5:**
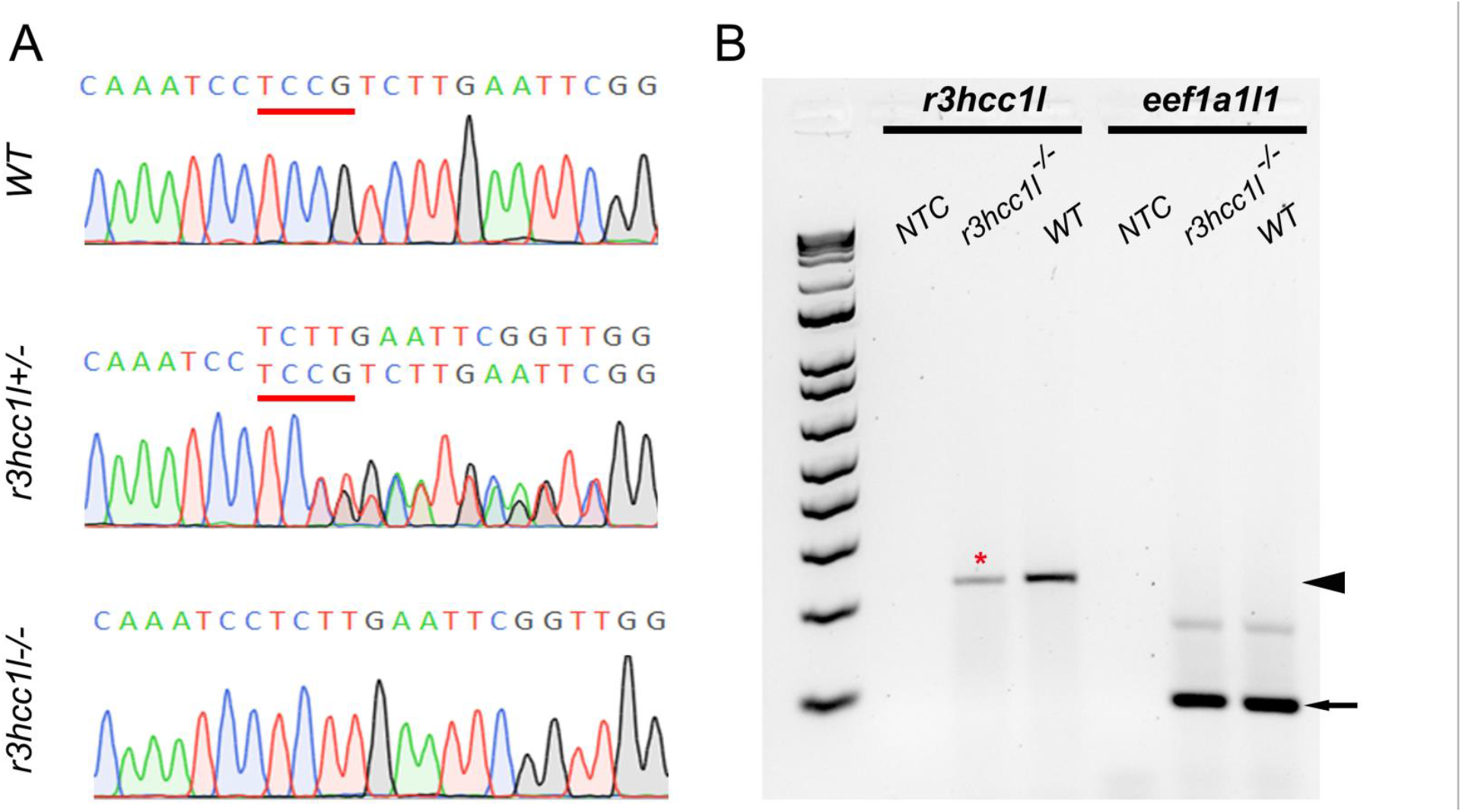
(A) Chromatograms showing DNA sequences for wild-type (top), heterozygous (middle), and homozygous (bottom) *r3hcc1l* mutants (in reverse direction). (B) Gel electrophoresis of RT-PCR products generated with *r3hcc1l* (black arrowhead; lanes 2-4) and *eef1a1l1* (black arrow; lanes 5-7) specific primers. Lane 1 – 1kb plus DNA ladder (Thermo Fisher Scientific); Lane 2 - no template control (NTC); Lane 3 - PCR product (231 bp) amplified from RNA extracted from homozygous *r3hcc1l*^*c*.*623-626del*^ embryos (marked with red asterisk); Lane 4 - PCR product (235 bp) amplified from RNA extracted from wild-type (WT) zebrafish embryos; Lane 5 - no template control (NTC); Lane 6 - PCR product (87 bp) amplified from RNA extracted from homozygous *r3hcc1l*^*c*.*623-626del*^ embryos; Lane 7 - PCR product (87 bp) amplified from RNA extracted from wild-type (WT) zebrafish embryos; Lanes 2-4 show products generated with *r3hcc1l*-specific primers, while Lanes 5-7- with *eef1a1l1*-specific primers. WT-wild-type.

Two founder (F0) pairs heterozygous for this mutation were selected and bred. F1 progeny had no gross abnormalities (n=160) when observed up to 5 dpf. At adult stages (>3 months) fish were also phenotypically normal (Figure 6). Genotypes of the F1 adults matched the expected Mendelian ratios (n=65 [19 homozygotes: 29 heterozygotes: 17 wild type], χ^2^=0.877, df=2, p=0.645) showing that homozygous mutants have normal survival. To rule out a potential protective role of maternal transcripts, a female homozygous fish was crossed with a male heterozygous fish. Genotypes of these F2 adults also matched the expected Mendelian ratios (n=13 [6 homozygous: 7 heterozygous] χ^2^=0.077, df=1, p=0.781) providing further evidence that homozygous mutants have normal viability. Finally, crosses between homozygous parents produced morphologically normal offspring that survived to adulthood at similar rates to wild-type animals.

**Figure 6:**
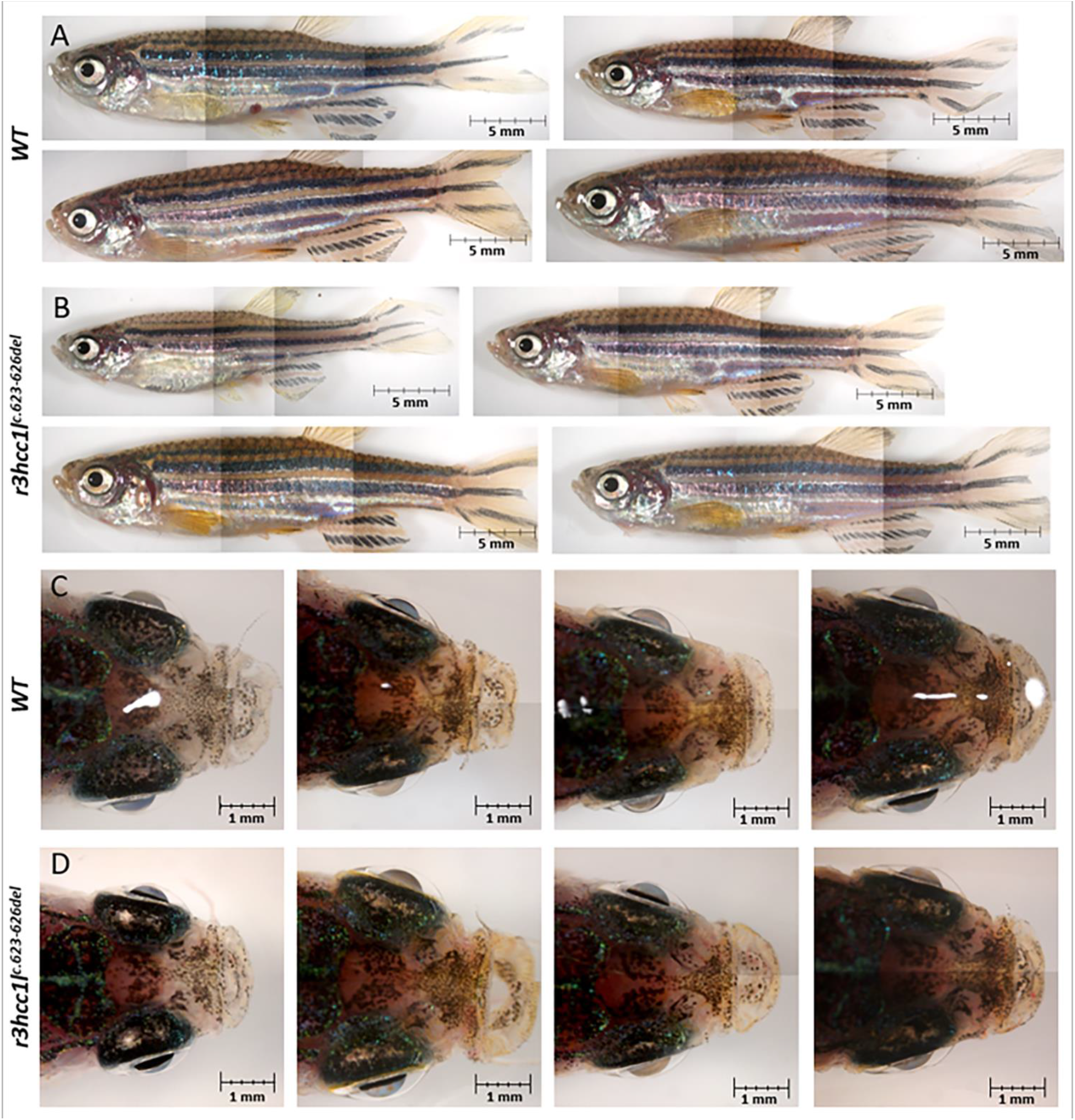
Lateral (A, B) and dorsal (C, D) views of wild-type (A, C) and homozygous *r3hcc1l*^*c*.*623-626del*^ (B, D) adult zebrafish (8 mpf). Please note overall normal morphology, including eye size and structures in the homozygous *r3hcc1l*^*c*.*623-626del*^ mutants. Size bars are indicated in the right bottom corner.

## Discussion

PA is a heterogeneous developmental disorder with many cases still unexplained. We identified compound heterozygous variants in *R3HCC1L* in one such case. *R3HCC1L* is a plausible candidate because of its predicted function and its expression in developing ocular tissues in vertebrates (zebrafish). Not much is known about the function of R3HCC1L, but it is predicted to be involved in post-translational gene regulation. R3HCC1L contains two conserved domains in its C-terminus that are shared by all known isoforms. One of them, an RRM, also known as the ribonucleoprotein (RNP) motif, is the most common RNA-binding motif and is present in proteins involved in RNA processing and transport such as heterogeneous nuclear RNPs, spliceosomal proteins, and poly(A)-binding proteins (SenGupta, 2013). The second conserved region, a coiled-coil domain (CCD), is likely a protein folding motif. These motifs are commonly found in fibrous proteins and transcription factors that are typically involved in the assembly of higher order protein structures such as transcription complexes, although other functions, including DNA binding, have also been identified (Lupas, 1996). Consistent with its predicted role as an RBP, *R3HCC1L* was found to localize subcellularly to the nucleoplasm and specifically the nuclear speckles ((Thul et al., 2017); Human Protein Atlas proteinatlas.org). Recent work investigating the contribution of RBPs to ocular development has shown that disruptions of several known RBPs (*Caprin2, Celf1/CELF1, Tdrd7/TDRD7*) lead to developmental defects of the eye such as PA, congenital cataract, and glaucoma in animal models and/or human patients (Aryal et al., 2020; Dash et al., 2015; Lachke et al., 2011). Of specific interest, *Caprin2* conditional knockout mouse lines exhibited two ocular phenotypes at variable penetrance: reduction in the lens nucleus or a persistent lenti-corneal stalk (Dash et al., 2015). This association of RBP genes with ocular developmental anomalies demonstrates a plausible mechanism for *R3HCC1L* in ocular development.

Since the vertebrate eye develops in an evolutionarily conserved manner, in terms of both developmental processes and underlying genetic networks, animal models can provide unique insight into the functional role of a specific factor. Mouse *R3hcc1l* is expressed broadly across all tested tissues including ocular structures ((Lattin et al., 2008; Wu et al., 2009) biogps.org). There are two known *R3hcc1l* transcripts in mice: NM_177464.4 encoding a protein corresponding to human isoform 2 (778 aa) and XM_006527182.4 encoding a protein corresponding to human isoform 3 (184 aa); mouse and human isoform 2 share the N-terminal region carrying the human variants of interest, thus showing higher overall conservation in comparison to zebrafish. While there are no published studies that examine a role of *R3hcc1l* during mouse development, an unpublished *R3hcc1l* gene deletion mouse line (*R3hcc1l*^*tm1b*^*(KOMP)*^*Wtsi*^) demonstrated ocular phenotypes, such as the persistence of the hyaloid vascular system and corneal opacity, but only in the female mice and only the persistence of the hyaloid vascular system reached the threshold for significance ((Groza et al., 2023) www.mousephenotype.org). Though preliminary, these results support the possibility that *R3HCC1L* deficiency may interfere with ocular development.

Studies in zebrafish presented here revealed embryonic expression of *r3hcc1l* at multiple sites including the developing lens, cornea, and hyaloid vasculature. CRISPR-Cas9 genome editing was used to generate a likely loss-of-function allele in *r3hcc1l*, c.623-626del p.(Thr208fs*39), that leads to truncation of the normal r3cch1l protein sequence and disrupts its two functional domains. However, the resulting heterozygous and homozygous animals did not show any opacities in the lens/cornea, as well as any other visible structural abnormalities in the eye or other systems, with normal survival of all genotypes to adulthood. This argues against a significant role for this gene in vertebrate embryonic development. However, it is possible that the function of *r3hcc1l* is not completely conserved with that of *R3HCC1L*, and/or that zebrafish have compensatory mechanisms not present in humans. The presence of multiple isoforms in humans and a single known transcript in zebrafish (encoding for a shorter isoform, lacking the exon affected in the human patient) should also be considered, highlighting the need for further studies into the possible broader spectrum of *r3hcc1l* isoforms in zebrafish, similar to humans and mice. If successful, then generation of *r3hcc1l* lines that more precisely mimic the identified human alleles can be attempted to further investigate their possible pathogenicity. Another approach to interrogate the roles of different human *R3HCC1L* isoforms and, specifically, the identified human variants, is to use an animal model in which orthologous transcripts have already been identified, such as the mouse. While it is important to consider the limitations of animal models in this case, it is also necessary to consider that the human variants in *R3HCC1L* are variants of unknown significance and may not have any role in the phenotype observed in this patient. While they were the strongest variants identified by exome sequencing, it may be necessary to pursue further analysis in this case through whole genome sequencing. In addition to further exploration of this case, continued analysis of similar patient cohorts, which may or may not lead to identification of new *R3HCC1L* variants of interest in families affected with PA or other congenital phenotypes, is still an important approach to further understand the contribution of the variants reported here.

## Acknowledgements

We are grateful to the family for participating in the study. This work was funded by the National Institutes of Health [R01 EY015518 to EVS].

